# Biogenesis of a young, 22-nt microRNA in Phaseoleae species by precursor-programmed uridylation

**DOI:** 10.1101/310920

**Authors:** Qili Fei, Yu Yu, Li Liu, Yu Zhang, Patricia Baldrich, Qing Dai, Xuemei Chen, Blake C. Meyers

**Affiliations:** Department of Plant & Soil Sciences, Delaware Biotechnology Institute, University of Delaware, Newark, Delaware 19711, USA; Donald Danforth Plant Science Center, St. Louis, Missouri 63132, USA; Department of Botany and Plant Sciences, Institute of Integrative Genome Biology, University of California, Riverside, California 92521, USA; Department of Chemistry and Institute for Biophysical Dynamics, The University of Chicago, Chicago, Illinois 60637, USA; Howard Hughes Medical Institute, The University of Chicago, Chicago, Illinois 60637, USA; Guangdong Provincial Key Laboratory for Plant Epigenetics, College of Life Sciences and Oceanography, Shenzhen University, Shenzhen, 518060, China; University of Missouri – Columbia, Division of Plant Sciences, 52 Agriculture Lab, Columbia, MO 65211, USA

**Author notes:** Present address: Department of Chemistry, University of Chicago, Chicago, IL 60637, USA. Corresponding author: Blake C. Meyers.

**Keywords:** microRNA evolution, soybean, 2’-O-methylation, uridylation, Phaseoleae, *NB-LRR*

## Abstract

Phased, secondary siRNAs (phasiRNAs) represent a class of small RNAs in plants generated via distinct biogenesis pathways, predominantly dependent on the activity of 22 nt miRNAs. Most 22 nt miRNAs are processed by DCL1 from miRNA precursors containing an asymmetric bulge, yielding a 22/21 nt miRNA/miRNA* duplex. Here we show that miR1510, a soybean miRNA capable of triggering phasiRNA production from numerous *NB-LRRs*, previously described as 21 nt in its mature form, primarily accumulates as a 22 nt isoform via monouridylation. We demonstrate that in Arabidopsis, this uridylation is performed by HESO1. Biochemical experiments showed that the 3’ terminus of miR1510 is only partially 2’-O-methylated, because of the terminal mispairing in the miR1510/miR1510* duplex that inhibits HEN1 activity in soybean. miR1510 emerged in the Phaseoleae ~41 to 42 MYA with a conserved precursor structure yielding a 22 nt monouridylated form, yet a variant in mung bean is processed directly in a 22 nt mature form. This analysis of miR1510 yields two observations: (1) plants can utilize post-processing modification to generate abundant 22 nt miRNA isoforms to more efficiently regulate target mRNA abundances; (2) comparative analysis demonstrates an example of selective optimization of precursor processing of a young plant miRNA.

## INTRODUCTION

Plant miRNAs are capable of triggering phased, secondary siRNAs (phasiRNAs) from long noncoding RNAs (lncRNA) or mRNAs (1). These phasiRNAs participate in both plant development and immunity. A number of 22 nt miRNAs, such as miR482/2118, miR1507, miR2109, miR5300, and miR6019, trigger phasiRNAs from the *nucleotide-binding leucine-rich repeat* (*NB-LRR*) gene family, which constitutes the majority of plant disease resistance (R) genes (12–15). *NB-LRR-* derived phasiRNAs have been confirmed to reinforce the efficacy of these 22 nt miRNA triggers in *NB-LRR* suppression. For instance, 22 nt miR9863 targets *Mla* transcripts triggering phasiRNAs, which, in concert with miR9863, represses *Mla* in barley (16). Consistent with this study on disease resistance, another report demonstrated that more widespread and efficient silencing was observed for a 22 nt artificial miRNA (amiRNA), relative to a 21 nt version, because of the generation of phasiRNAs (17). Consequently, it is postulated that NB-LRR-derived phasiRNAs act as an essential layer to fine-tune *R* gene expression (18). miR1510 is a notable miRNA in this same class; it’s a legume-specific miRNA that is the predominant trigger of phasiRNAs from *NB-LRRs* in soybean, yielding abundant phasiRNAs from its targets (19). Yet, given its annotated length as 21 nt, it has been unclear why this miRNA processed has such a substantial activity as a trigger of phasiRNAs.

Plant small RNAs, including both miRNAs and siRNAs, are extensively subject to the modification of 3’ terminal 2’-O-methylation by the methyltransferase HUA ENHANCER 1 (HEN1) (20, 21), preventing small RNAs from 3’ uridylation and subsequent degradation (22). In a *hen1* mutant background, and thus in the absence of 3’ methylation protection, miRNAs tend to be truncated from the 3’ end prior to uridylation mediated by different nucleotidyltransferases, such as HEN1 SUPPRESSOR 1 (HESO1) and UTP:RNA URIDYLYLTRANSFERASE 1 (URT1) (23–27). Consequently, miRNA abundances are generally reduced in the *hen1* mutant, resulting in pleotropic developmental defects (28), while different miRNAs display distinct patterns of truncation and tailing (23, 24). One unusual gain-of-function in the *hen1* mutant background was observed: miR171a triggers phasiRNA production from target transcripts, because the typically 21 nt mature miRNA is abundantly tailed to 22 nt by URT1 in absence of 2’-O-methylation (24, 25). This observation supports that the 22 nt length of miRNAs is important for phasiRNA production.

Here we show that in wildtype soybean, 21 nt miR1510 is partially methylated and subsequently uridylated to 22 nt by HESO1, likely bestowing on miR1510 the ability to trigger phasiRNA production from target transcripts. We found that the mismatch adjacent to the 2 nt 3’ overhang in the miR1510/miR1510* duplex inhibits HEN1 activity *in vitro*, resulting in its 3’ monouridylation by HESO1. Interestingly, the position of the mismatch is conserved across the Phaseoleae tribe of legume species, and high levels of uridylated miR1510 in 22 nt form were also observed in other Phaseoleae species, including common bean and pigeon pea. Therefore, we propose that the Phaseoleae have evolved to employ this mechanism to generate a 22 nt miRNA and its consequential phasiRNAs to fine-tune *R* gene expression.

## RESULTS

### In soybean, 21 nt miR1510 is predominantly uridylated to 22 nt

miR1510 targets transcripts of over 100 *NB-LRRs* genes in soybean, far more than any other miRNA, triggering abundant phasiRNAs from their transcripts (19, 29). The mature miRNA is generated from two *MIR1510* loci in the soybean genome, copies that likely originated from the genome duplication during soybean evolution (30) (Fig. 1A). Based on these analyses of the precursor, miR1510 is processed into a 21 nt mature miRNA (Fig. S1), likely by DICER-LIKE1 (DCL1) as in other species; yet, based on numerous previous studies, a length of 22 nt is typically required for phasiRNA biogenesis. We therefore investigated why this 21 nt miRNA is capable of triggering phasiRNAs. Unexpectedly, a search of miR1510 reads in small RNA sequencing data showed that the most abundant form of miR1510 is a 22 nt isoform (Fig. 1A). This 22 nt miR1510 does not map to the soybean genome, because of the additional 22^nd^ nucleotide (the 3’ end), a uridine (Fig. 1A), perhaps explaining why it was previously overlooked. We assessed whether this isoform could have been generated from a precursor missing (i.e. in a gap) in the current soybean genome assembly. To do so, we examined RNA-seq data that would include pri-miRNA transcripts. However, we found no RNA-seq reads containing the 22 nt isoform of miR1510, suggesting that it is not generated from the genome; instead, the ‘U’ at the 22^nd^ position is more likely the result of uridylation, possibly by a member of the nucleotidyltransferase family.

**Figure 1.**
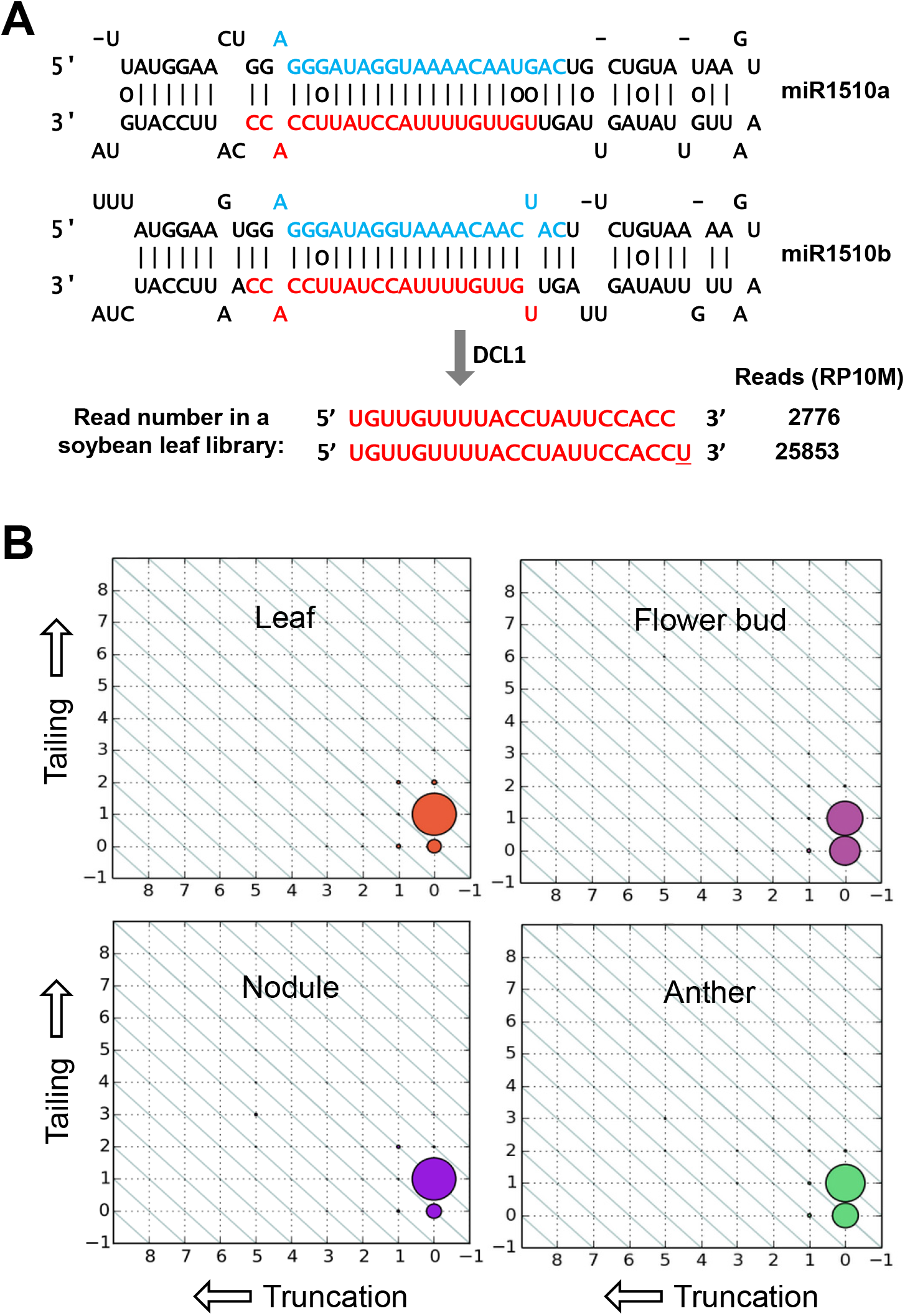
miR1510 is predominantly tailed to 22-nt in wildtype soybean. A. Precursor sequences and predicted secondary structures of miR1510a and miR1510b precursors in soybean. The miRNA strand is indicated in red, while the miRNA* strand is in blue. Sequencing data of a small RNA library from the leaf tissue is shown below the arrow. Normalized read numbers are shown on the right. B. Truncation and tailing status of miR1510 in different tissues of wildtype soybean.The status of 3’ truncation and/or 3’ tailing for miR1510 is represented by a two-dimensional matrix. The sizes of the dots indicate the relative abundance of different isoforms of miR1510. The x axis represents the extent of 3’ truncation, with the canonical, non-truncated 21-nt length at the 0 position (at right); the y axis represents 3’ tailing, with the canonical, non-tailed 21-nt length at the 0 position (at bottom). Dots on the diagonals represent reads of the same length but with different amounts of truncation/tailing. Soybean small RNA sequencing data were from GSE58779 (Leaf - GSM1419325, Flower bud: GSM1419321, Anther: GSM1419316, Nodule: GSM1419376).

We next examined how broadly the monouridylated form of miR1510 exists in different tissues of soybean. We employed a previously described method for analysis of truncation and tailing of miRNAs (31). We analyzed the published data that comprises an atlas of soybean small RNAs (29); we observed that as with the leaf tissue miR1510 is uridylated to 22 nt in other tissues, including nodule, flower, and anther, although the degree of uridylation varies (Fig. 1B). This result indicated that the 22 nt form of miR1510 accumulates abundantly in a variety of tissues of wildtype soybean, consistent with its role as a dominant trigger of phasiRNAs from *NB-LRR* targets. In contrast, other miRNAs in wildtype soybean, such as miR172, miR396, miR398 and miR482, have barely measurable levels of truncated or tailed forms (Fig. S2), suggesting that miR1510 is unique among soybean miRNAs in its tailing.

### Non-species-specific uridylation of miR1510 in plants

Considering that miR1510 was the only miRNA for which we observed significant uridylation in soybean, we hypothesized that this miRNA may have attributes, such as a specific precursor structure, that facilitate the uridylation, and thus perhaps it would be uridylated in other plant species. To this end, we transformed both *MIR1510a* and *MIR1510b* into Arabidopsis to make stable transgenic lines, and also transiently expressed both precursors in *Nicotiana benthamiana*. Small RNA gel blotting showed that *MIR1510a* generated abundant 22 nt miRNAs when expressed in Arabidopsis, as in soybean (Fig. S3A). However, mature miR1510 was not detected from *MIR1510b* transformants, likely because it was not processed by DCL1 in Arabidopsis, as RT-PCR experiments verified the expression of *MIR1510b* in Arabidopsis (Fig. S3B). We speculate that the processing of *MIR1510b* by Arabidopsis DCL1 is inhibited by two mismatches at the sites that if cut would release the miR1510b/miR1510b* duplex (Fig. 1A), which is due to the divergence of DCL1 among species. In the *N. benthamiana* transient expression assays, as in soybean and Arabidopsis, considerable amount of 22 nt miR1510 was also detected, suggesting that miR1510 is monouridylated in both Arabidopsis and tobacco.

To rule out the possibility that the 22 nt isoform of miR1510 was generated via imprecise processing by DCL1, we sequenced the small RNAs from the Arabidopsis leaf sample transformed with *MIR1510a*. Sequencing data showed that 22 nt miR1510 was indeed generated by uridylation, and its abundance was > 2 fold higher than the 21 nt form (Fig. S3C). Collectively, our data showed that the uridylation of miR1510 is non-species-specific, and we inferred that its precursors encode features that trigger monouridylation of the mature miRNA.

### miR1510 is uridylated by HESO1, but not URT1

Two nucleotidyltransferases in Arabidopsis, including HESO1 and URT1, have a demonstrated ability to uridylate the 3’ terminus of miRNAs (23, 25–27). These two nucleotidyltransferases show different preferences in miRNA substrates *in vitro*, and their substrate preference is largely determined by the 3’ terminal nucleotide of the miRNA (25). We subsequently explored whether the 22^nd^ ‘U’ of miR1510 is added by HESO1, URT1, or possibly some other nucleotidyltransferase yet to be characterized. To investigate this, *MIR1510a* was transformed separately into *heso1-1* and *urt1-1* homozygous mutant backgrounds, and we sequenced the small RNAs from the leaf tissue of the screened T1 plants. Sequencing data showed that the 22 nt form of miR1510 was substantially decreased in a *heso1-1*, but not a *urt1-1* mutant background (Fig. 2). This genetic analysis showed that HESO1 is responsible for the monouridylation of miR1510 in Arabidopsis *in vivo*, and potentially by its orthologs in soybean and tobacco.

**Figure 2.**
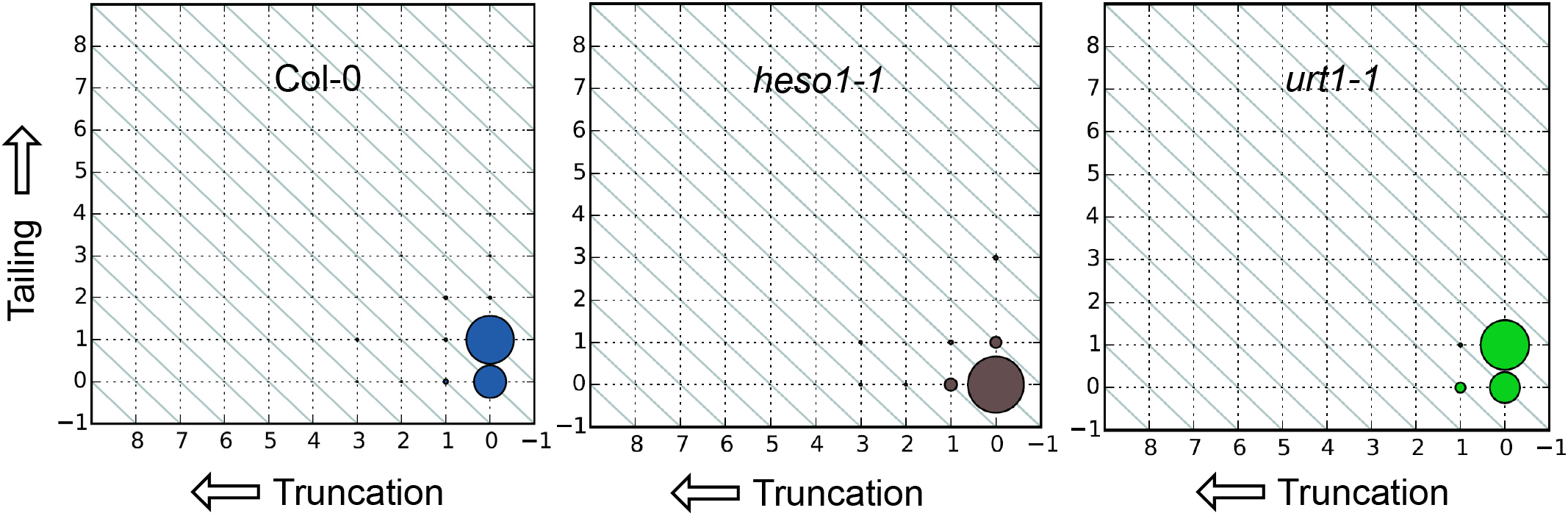
Truncation and tailing analysis of miR1510 expressed in different genotypes of Arabidopsis. Small RNA libraries were prepared from the leaf tissue of Arabidopsis, expressing the miR1510a precursor from a stable transgene as T1 lines in wildtype (Col-0), or *heso1-1* or *urt1-1* homozygous mutants. Reads corresponding to miR1510 were identified in each library and analyzed for 3’ truncation and tailing, and plotted for each genotype, as indicated; the plots are interpreted as described in Figure 1.

### miR1510 is partially methylated in soybean

Plant miRNAs are extensively methylated at the 3’ terminus by HEN1 to avoid uridylation and subsequent miRNA turnover (22). The uridylation of miR1510 suggests that this miRNA is possibly partially methylated at the 3’ terminus. To test this, β-elimination was employed to assess the methylation status of miR1510 in soybean. In parallel with the oxidative treatment of NaIO4 for miR1510, we tested miR166 using total RNA samples from the Arabidopsis *hen1-8* loss-of-function mutants, in which miRNAs are predominantly unprotected by 2’-O-methylation at the 3’ terminus (22). The treated samples showed a shifted band running ~2 nt faster than the untreated miR166 band in the *hen1-8* background (Fig. S4), confirming that the β-elimination treatment was successful. For soybean miR1510, we found that an additional 19 nt band, although weak, appeared after the treatment, revealed by small RNA gel blotting (Fig. 3). This result suggested that the 21 nt isoform of miR1510 is partially methylated Unexpectedly, β-elimination did not generate an increased intensity of the 20 nt band for miR1510 in treated samples (Fig. 3), indicating that the 22 nt monouridylated miR1510 variant is protected by 2’-O-methylation, presumably by the methyltransferase activity of HEN1. This result also suggests that the methylation after uridylation occurs to AGO-bound miRNAs, because of observations that uridylated miRNAs are bound by Argonaute proteins (24, 25, 32). Previous studies in *Drosophila melanogaster* showed that its HEN1 homolog, DmHen1, methylates PIWI-bound piRNAs and Ago2-bound siRNAs (33, 34), implying that HEN1 may methylate AGO-bound small RNAs in plants.

**Figure 3.**
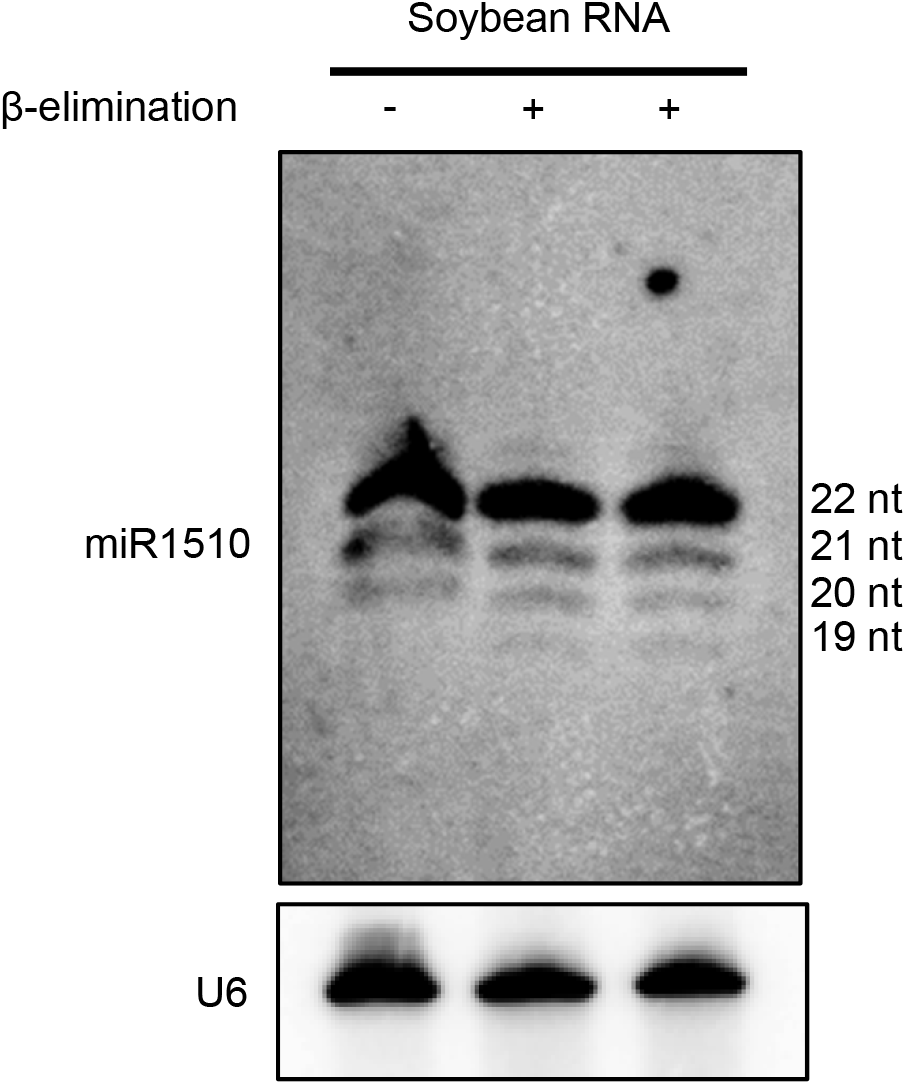
Partial 3’-terminal 2’-O-methylation of soybean miR1510, revealed by β-elimination. Untreated RNA is loaded on the left (−), while two replicate, treated samples are on the right (+). The appearance of a 19 nt band in the two treated samples but not the untreated control is consistent with partial methylation of miR1510. All samples were RNA extracted from wildtype soybean leaves.

### miR1510 is a young miRNA, specific to the Phaseoleae tribe of legumes

The unique features of miR1510 led us to investigate the evolution of this miRNA in plants. Using *MIR1510a/b* precursor sequences from soybean to search in the currently-available, sequenced genomes of legume species, we found that only a few legume species contain precursor sequences, including wild soybean (*Glycine soja*), common bean (*Phaseolus vulgaris*), pigeon pea (*Cajanus cajan*), and mung bean (*Vigna radiata*). A further check of these species revealed that they all belong to the Phaseoleae tribe of the Papilionoideae subfamily, which diverged ~41 to 42 million years ago (MYA) (Fig. 4A) (35). In miRBase (release 21), we observed that records exist for miR1510a/b in *Medicago truncatula*, and therefore we compared their precursor sequences with those in soybean. Sequence alignment showed that the miR1510 precursor sequences and even the mature miRNAs in Medicago and soybean differ substantially, indicating that miR1510 in Medicago has a different origin and is essentially a distinct miRNA (Fig. S5). Compared with other NB-LRR-targeting miRNAs such as the ancient miR482/2118 superfamily (36, 37), miR1510 is an evolutionarily young miRNA, specific to the Phaseoleae tribe of legumes.

**Figure 4.**
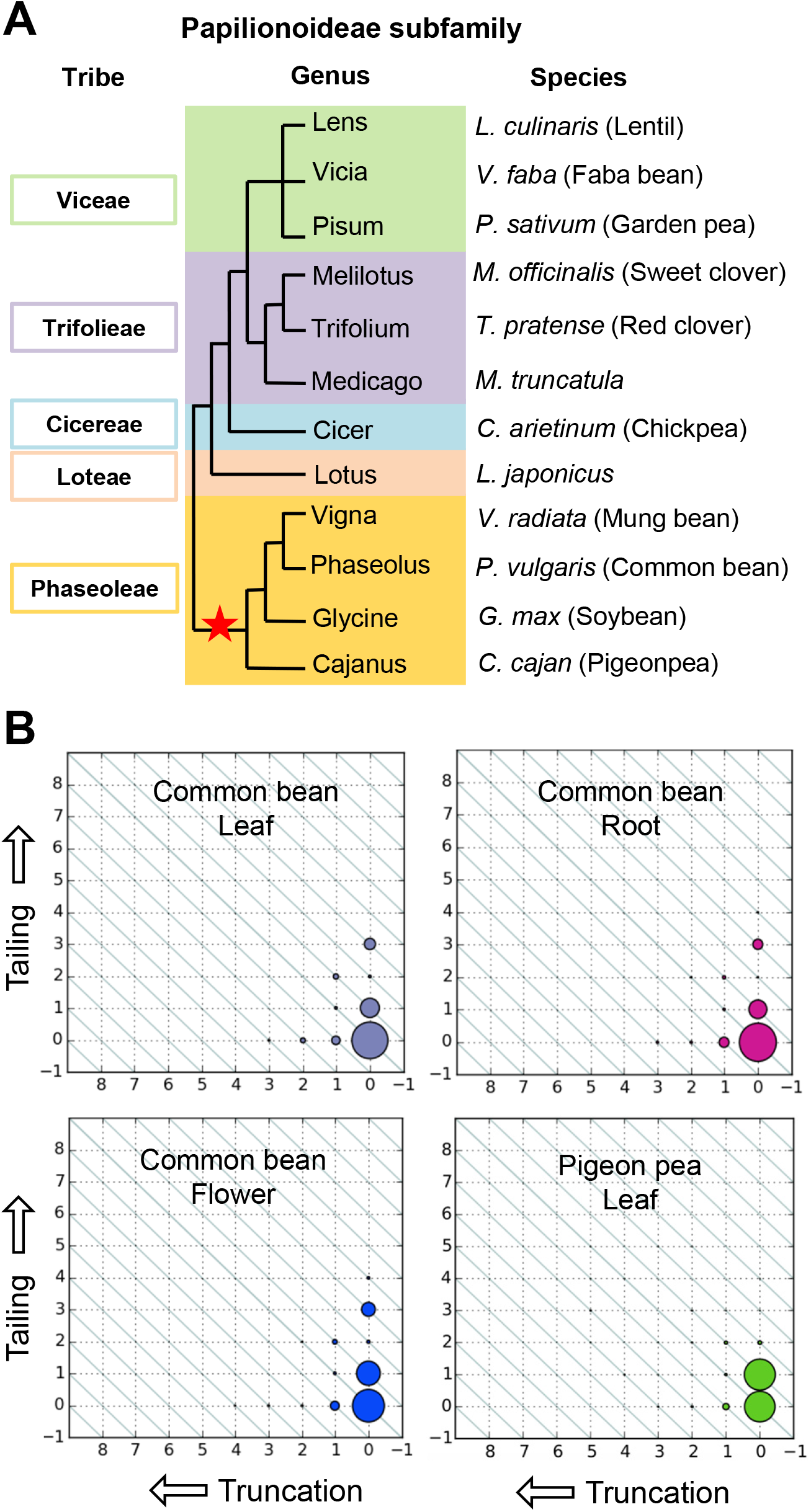
Monouridylation of miR1510 is exclusive to the Phaseoleae species. A. Phylogeny of the Papilionoideae subfamily of legumes. The Papilionoideae subfamily consists of five tribes, each of which contains one to multiple genera. Representative species are indicated at right. Red star indicates that miR1510 is present exclusively in the Phaseoleae tribe. The phylogenetic tree is adapted from Choi et al., 2004. B. Truncation and tailing analysis of miR1510 in common bean and pigeon pea; tissues are as indicated below the plant type in each plot. A small RNA library of pigeon pea was prepared from leaf tissue. Common bean data were from the published dataset GSE67409. The plots are interpreted as described in Figure 1.

### Abundant 22 nt form of miR1510 in Phaseoleae species

The conservation of miR1510 in Phaseoleae species gave rise to the question whether this miRNA is commonly 22 nt because of uridylation. We examined this by checking both published small RNA sequencing data for common bean and preparing small RNA libraries from the leaf tissue for species without published datasets, including pigeon pea and mung bean. Truncation and tailing analysis in common bean indicated that a considerable proportion of miR1510 is uridylated to 22 nt (Fig. 4B). In contrast, pigeon pea has a larger fraction (~50%) of monouridylated miR1510 compared with common bean (Fig. 4B). The variation of the levels of 22 nt miR1510 in diverse tissues and species possibly result from differences in levels of miR1510 and HESO1 across tissues, or from altered activities of HEN1 in these species.

The processing of miR1510 in mung bean seems different from that other Phaseoeae species. Small RNA sequencing data showed that the mature miR1510 in mung bean contains a 2 to 3 nucleotides shift compared with other species. Interestingly, in mung bean, miR1510 is likely processed directly into 22 nt by DCL1, because the 21 nt miR1510 read was absent in the small RNA data for mung bean. In addition, the 22 nt forms of both miR1510 and miR1510* form a duplex with a 3’ single nucleotide overhang, which is atypical for plant miRNAs (Fig. S6). Therefore, miR1510 in mung bean is 22 nt in length, which derives from the direct processing of DCL1, without uridylation. Taken together, the 22 nt forms of miR1510 are universally abundant in Phaseoleae species and predominantly generated by uridylation, although miR1510 is likely directly processed into 22 nt by DCL1 in mung bean.

A previous study integrating small RNA and PARE data reported that miR1510 triggers phasiRNAs from 20 *NB-LRRs* in soybean (29). We reexamined the data and checked how the 22 nt miR1510 pairs with these targets. Among the 20 reported genes, 11 passed the default filter of psRNATarget (38), which assesses base pairing between miRNAs and target mRNAs. Eight out of the 11 targets of 22 nt miR1510 showed 3’ terminal pairing, including both A:U and G:U wobble (Table S1). This result is consistent with previous reports that the terminal pairing of a 22 nt miRNA is important for phasiRNA production (15, 39), suggesting that 22 nt miR1510 might be under selection for the capacity to trigger phasiRNAs.

### Terminal Mispairing in the miR1510/miR1510* Duplex Inhibits HEN1 Methyltransferase Activity *in vitro*

Knowing that 21 nt miR1510 is incompletely methylated at its 3’ terminus, we speculated that the activity of HEN1 might be inhibited, resulting in uridylation. Previous studies showed that HEN1 recognizes miRNA/miRNA* duplexes and deposits a methyl group at the 3’ terminus (20, 21). Thus, we reasoned that miR1510/miR1510* structure might hamper the activity of HEN1. In examining the secondary structure of the miRNA precursors, we found that all secondary structures predicted for miR1510 precursors contain a mismatch at the 5’ terminal nucleotide of miR1510* (Fig. S7). This mismatch would introduce a terminal mispairing in the miR1510/miR1510* duplex after DCL1 processing that could potentially interfere with HEN1 activity.

To further test if the terminal mispairing of the miR1510/miR1510* duplex inhibits HEN1 activity, we conducted *in vitro* assays. We annealed synthetic RNA oligonucleotides of miR1510 (oligo #1) and miR1510* (oligo #2); one variant included a mutated miR1510* (oligo #3) to form a terminal pairing structure with miR1510 (Fig. 5A). These two different types (“mismatch” and “match”) of duplexes were then incubated with purified recombinant GST-HEN1 protein (Fig. S8A) in the presence of S-adenosyl methionine (SAM). We digested the RNA into single nucleotides using nuclease P1 followed by dephosphorylation with alkaline phosphatase, and then analyzed the ratios of 2’-O-methylated cytidine (Cm) to guanosine (G) by liquid chromatography with tandem mass spectrometry (LC-MS/MS). We found that the HEN1 activity was considerably lower (~50%) for the “mismatch” form of the duplex (Fig. 5B), which is the natural soybean miR1510/miR1510* duplex structure, than the mutated “match” form, indicating that the terminal mismatch can indeed inhibit HEN1 methyltransferase activity. In addition, we observed that HEN1 activity is extremely low but detectable for single-stranded RNA oligos (Fig. S8B), consistent with prior work (20).

**Figure 5.**
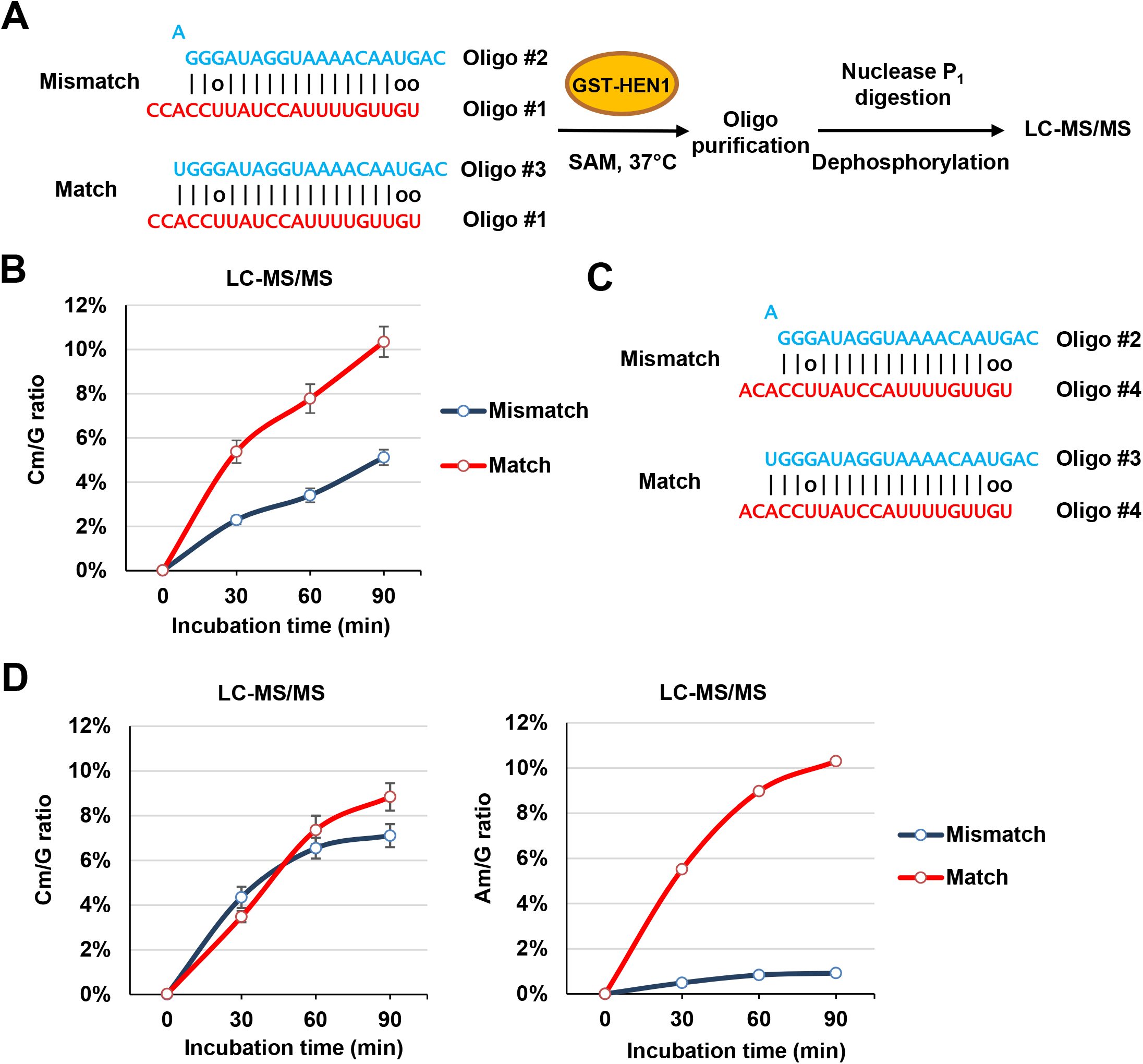
HEN1 methyltransferase activity assays. A. Work flow of the HEN1 methyltransferase activity assays. RNA oligos were annealed as indicated to form a “mismatch” or “match” structure. The annealed duplexes were incubated with purified GST-HEN1 protein, in the presence of SAM at 37° C. Oligos were purified after the reaction, and then digested into single nucleotides by nuclease P1, followed by dephosphorylation by FastAP before LC-MS/MS. B. Ratio of 3’-O-methylated C (Cm) to G. Levels of Cm to G were quantified by LC-MS/MS for two forms of duplexes after incubation with GST-HEN1 at 0, 30, 60 and 90 minutes. Means ± s.d. obtained from three individual HPLC injections. C. Oligo #4 annealed with Oligo #2 and #3 form “mismatch” and “match” duplex structures. Oligo #4 has a 3’ terminal A instead of C to distinguish the methylation strand. D. Ratios of Cm (left panel) and Am (right panel) to G were quantified by LC-MS/MS. Means ± s.d. obtained from three individual HPLC injections.

Because both miR1510 and miR1510* possess cytidines at the 3’ terminal, it was indistinguishable from which strand the Cm originated. We therefore investigated whether HEN1 methyltransferase activity was equally inhibited by the terminal mismatch at both strands because of (a) possible weakened HEN1 binding, or (b) reduced methyltransferase activity exclusively at the miR1510 strand. This experiment utilized oligo #4, which incorporated a terminal adenosine instead of cytidine (Figure 5C). LC-MS/MS results showed similar ratios of Cm/G for both “mismatch” and “match” forms of duplexes at all examined time points (Figure 5D, left panel), demonstrating that the methylation of the miR1510* strand was not disrupted by the mismatch on the other terminus. In contrast, Am/G ratio was consistently much lower (~10%) in the “mismatch” form (Figure 5D, right panel), demonstrating that the natural terminal mispairing largely reduced the catalytic rate of HEN1 on the miR1510 strand, consistent with the gel blot result demonstrating that 21 nt miR1510 is partially methylated (Fig. 3). Moreover, our experiments demonstrated that the terminal mismatch specifically inhibits HEN1 methyltransferase activity at the corresponding duplex overhang, instead of weakening HEN1 binding to the RNA duplex to inhibit methylation at both strands.

## DISCUSSION

Soybean is a paleopolyploid and experienced two rounds of genome duplications about 59 and 13 million years ago (30), resulting in over 500 *NB-LRRs* in the genome (19). miR1510 targets *NB-LRRs* in soybean, and with >100 predicted targets is the major miRNA targeting this family, triggering the production of phasiRNAs from transcripts of many genes (19). The 21 nt isoform of miR1510 was believed to be the trigger of phasiRNAs in soybean; however, this is inconsistent with previous studies showing that the 22 nt length of miRNAs is required for phasiRNA production (10, 11). We found that soybean contains high levels of a 22 nt isoform of miR1510, a length that is able to trigger phasiRNA production. The 22 nt isoform was largely missed in previous analyses, probably because it does not map to the soybean genome because of the additional, 22^nd^ nucleotide, a uracil (Fig. 1).

We investigated the biogenesis of the 22 nt form of miR1510, and found that it results from monouridylation. β-elimination showed that the 21 nt, processed form of miR1510 is partially methylated, making it susceptible to 3’ modification, and its subsequent uridylation (Fig. 3). Genetic analysis by transforming *MIR1510a* to both *hesol* and *urtl* mutant background revealed that HESO1, but not URT1, is responsible for the monouridylation of miR1510 *in vivo*. We found that miR1510 is generally uridylated among Phaseoleae species, and that the secondary structures of miR1510 precursors in different species generate a terminal mispairing in the miR1510/miR1510* duplex that inhibits HEN1 methyltransferase activity, resulting in HESO1-mediated uridylation. Indeed, HEN1 methyltransferase activity assays *in vitro* showed that the innate terminal mispairing of miR1510/miR1510* predominantly inhibits HEN1 methyltransferase activity at the miR1510 3’ terminus, but not miR1510*. However, the methyltransferase activity was partially (10%) maintained for miR1510, consistent with the endogenous methylation levels of 21 nt miR1510 in soybean (Fig. 3). We have integrated these observations in a model in which the precursors of miR1510 in Phaseoleae species contain this mispairing at a conserved position, yielding a 3’ terminal mispairing in miR1510/miR1510* duplexes after DCL1 processing (Fig. 6). The terminal mispairing in a duplex inhibits HEN1 methyltransferase activity for miR1510, but not the miR1510* strand. The methylation, Argonaute loading, and uridylation steps are tightly coordinated. miR1510 lacking 2’-O-methylation or those undergoing delayed methylation by HEN1 are loaded to AGOs, and thereby uridylated by HESO1, converting miR1510 to 22 nt, as prior work showed that HESO1 uridylates AGO-bound miRNAs *in vitro* (25). The AGO-bound 22 nt miR1510 resulting from uridylation might undergo methylation by HEN1, maintaining its stability, because we observed that the 22 nt isoform of miR1510 is fully methylated (Fig. 3). Therefore, the pathway that we demonstrate here is a coordinated process, coupling miRNA biogenesis and modifications.

**Figure 6.**
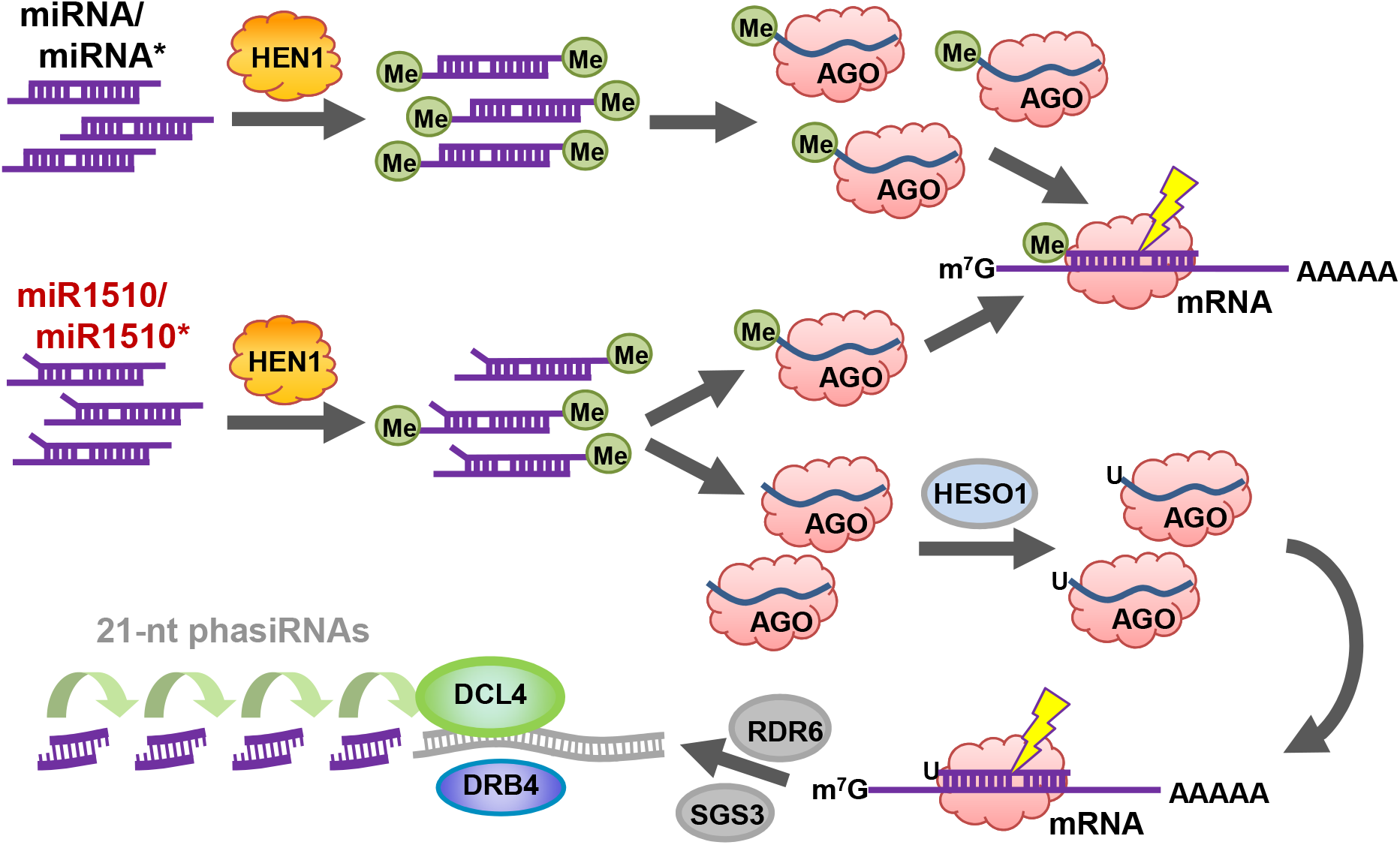
A model for the biogenesis and function of the 22 nt isoform of miR1510. Compared with other canonical miRNA/miRNA* duplexes, miR1510/miR1510* contains a terminal mispairing, which inhibits HEN1 methyltransferase activity on the miR1510 strand, but not the miR1510* strand. Typically, methylated miRNAs are loaded into AGOs for downstream RNA silencing. Unmethylated miR1510 is still loaded into AGOs, followed by 3’ terminal uridylation by HESO1. The AGO-bound uridylated miR1510 may be further methylated, possibly by HEN1 (not shown). The 22 nt miR1510 then triggers phasiRNA production by targeting RNA transcripts, with phasiRNA production requiring the involvement of RDR6, SGS3 and DCL4.

Systematic characterization of *MIR1510* genes in sequenced plant genomes showed that miR1510 is lineage-specific to the Phaseoleae tribe of legumes, which diverged ~41 to 42 MYA among the Papilionoideae subfamily. Thus, miR1510 is a relatively young miRNA in evolutionary terms, and variation in its processing/maturation steps may reflect a process of ‘optimization’. While most tested Phaseoleae species, including soybean, common bean, and pigeon pea, display the same mechanism to generate the abundant 22 nt isoform of miR1510, the biogenesis of miR1510 in mung bean seems distinct. Sequencing data showed that miR1510 in mung bean (“vra-miR1510”) is processed directly to 22 nt due to a shift in cleavage sites, although the mispairing at the same position was still retained in mung bean (Fig. S6). The direct processing of vra-miR1510 to 22 nt is supported by the absence of detectable levels of its 21 nt isoform, which is found at considerable levels as the un-uridylated form in other Phaseoleae species we examined. The recent evolutionary origin of miR1510 may explain its plasticity in DCL1 processing observed across Phaseoleae species, specifically the loss of a requirement for uridylation in mung bean to produce the 22 nt isoform.

miRNAs are important regulators of plant *NB-LRR* disease resistance genes. Evolutionary analysis demonstrates that *MIRNA* genes are occasionally generated from their target genes as a result of small-scale genome rearrangements such as duplications forming inverted repeats (44). Expanded counts of *NB-LRR* in plant genomes may be balanced by the emergence of miRNAs that target them (37), perhaps to minimize fitness costs and to avoid autoimmune responses (1). These miRNAs are often 22 nt, a length endowed with the property of triggering phasiRNAs, thereby reinforcing the efficacy of silencing via the secondary siRNAs which may have additional targets and mobility, potentially functioning to maintain a basal level of *NB-LRR* gene expression (16, 17). The 22 nt isoform of miR1510 found in the Phaseoleae is special in terms of its biogenesis, requiring monouridylation to achieve its length. Considering that this pathway involves multiple steps and that a considerable abundance of the 21 nt isoform remains detectable (i.e. the pathway is inefficient), an evolutionary advance would be the direct processing by DCL1 of *MIR1510* precursors into a 22 nt isoform – exactly as observed in mung bean. Therefore, we believe that via comparative genomic analysis, we have captured evidence of selection optimizing the processing of a plant miRNA precursor.

## EXPERIMENTAL PROCEDURES

Plant materials, vector construction and transformation, β-elimination, RT-PCR, small RNA sequencing and bioinformatics analysis, HEN1 purification and *in vitro* methyltransferase activity assays, LC-MS/MS, and primers and probes used in this study are described in SI Materials and Methods.

## AUTHOR CONTRIBUTIONS

B.C.M. and X.C. conceived the experiments. Q.F., Y.Y., L.L., P.B., and Q.D. performed experiments. Q.F. and Y.Z. conducted bioinformatics analyses. Q.F. and B.C.M. wrote the manuscript, with contributions from all authors.

## ACKNOWLEDGEMENTS

We thank Mayumi Nakano, S. Deepthi Ramachandruni, and Parth Patel for assistance with data handling and data visualization. We thank Dr. Scott Jackson for sharing the mung bean seeds. This work was supported by the US National Science Foundation award #1257869 from the Division of Integrative Organismal Systems (IOS) in the Meyers lab. Research in the Chen lab is supported by grants CA-R-BPS-5084-H and 2010-04209 from USDA-NIFA.

## Supplementary Information

### Supplementary Materials and Methods

#### Plant materials

*Glycine max* (Williams 82), *Arabidopsis thaliana* (Col-0), *Nicotiana benthamiana*, and *Vigna radiata* (VC1973A) plants were grown in a growth chamber or greenhouse with a light cycle of 16 h light / 8 h dark, at 22 to 23 °C. *Cajanus cajan* plants were grown in a greenhouse with a light cycle of 16 h light / 8 h dark, at 24 to 28 °C. The leaf tissues used for RNA extraction were all collected from 2-week old plants. The Arabidopsis mutant lines *heso1-1* and *urt1-1* were described previously (1).

#### Vector construction

MIR1510a/b genes were synthesized, amplified by PCR, inserted into Gateway pENTR Vectors (Invitrogen, Carlsbad, CA), and confirmed by Sanger sequencing. LR reactions were performed to transfer the *MIRNA* genes into binary vector pGWB402 containing the CaMV 35S promoter upstream of the cloning site. These constructs were used for Arabidopsis (Col-0) transformation. Vector pGWB502, which contains a hygromycin resistance gene, was used to transform Arabidopsis mutant lines. Primer sequences used for PCR amplification are listed in Table S2.

#### Plant transformation

Transient expression experiments in N. benthamiana were performed according to procedures described previously (Sparkes et al., 2006). Briefly, Agrobacterium tumefaciens carrying binary vectors was cultured at overnight at 28°C, and resuspended in infiltration buffer at OD600 = 0.4. Infiltrated leaves were collected 48 hours after infiltration. For stable transformation of Arabidopsis, the floral dip method was employed to transform both wildtype and mutant lines using binary vectors to permit expression of *MIR1510a/b*. A. tumefaciens GV3101 was used for both transient expression assays in N. benthamiana and floral dip transformation in Arabidopsis, as described above. Floral dip transformation was conducted as previously described (2).

#### RNA extraction, β-elimination and RNA gel blot hybridization

Total RNA samples in this study were purified from plant materials using PureLink Plant RNA Reagent (Ambion, Foster City, CA). β-elimination was conducted according to the method previously described (3). Briefly, 50 μg total RNA was treated with sodium periodate (20 mM) in borax/boric acid buffer (0.06 M, pH 8.6) for 60 min at room temperature in dark, followed by the addition of 10 μl glycerol to stop the reaction, with continued incubation for 10 min. RNA was then precipitated and treated with borax/boric acid/NaOH (0.055 M, pH 9.5) at 45 °C for 90 min. The reactions were then desalted using G-25 Columns (GE Healthcare Bio-Sciences, Pittsburgh, PA). Twenty μg desalted RNA samples and untreated total RNA samples were loaded into 20% denaturing polyacrylamide gels for overnight electrophoresis using a Protean II vertical electrophoresis system (BioRad, Hercules, CA). RNA was then transferred to nylon membrane, and crosslinked using 1-ethyl-3-(3-dimethylaminopropyl)carbodiimide (4). DNA oligo probes were end-labeled with γ-32P-ATP using T4 Polynucleotide Kinase (New England BioLabs, Ipswich, MA). Arabidopsis U6 small nuclear RNA was used as a loading control. Phosphor screens were scanned using a Typhoon scanner (GE Healthcare BioSciences, Pittsburgh, PA) after overnight exposure with the hybridized membrane. Probes used in this study are listed in Table S2.

#### RT-PCR

One μg total RNA from Arabidopsis leaf samples were treated with DNaseI (New England BioLabs, Ipswich, MA), and reverse-transcribed by SuperScript III Reverse Transcriptase (Invitrogen, Carlsbad, CA). The reverse transcription product was then amplified via PCR using gene specific primers (Table S2), followed by electrophoresis on agarose gels.

#### Small RNA sequencing and data analysis

Total RNA samples were used to construct small RNA libraries using the TruSeq Small RNA Sample Preparation Kit (Illumina, Hayward, CA) according to the manufacturer’s manual. The small RNA libraries were sequenced using an Illumina HiSeq instrument at either the Delaware Biotechnology Institute (Newark, DE) or MOgene (St. Louis, MO).

Small RNA reads were trimmed to remove the adapters, and then mapped to the corresponding plant genomes using Bowtie (5). Detailed processing and read normalization was performed as previously described (6). Processed data were then used as input for small RNA truncation and tailing analysis using the online tool miTRATA (7). miRNA target predictions were performed using psRNATarget using default settings (8).

#### Protein expression and purification

Expression vector pGEX-2TK-HEN1 was transformed to competent E. coli strain BL21 (New England BioLabs, Ipswich, MA), and then cultured in LB medium at 37 °C until OD600 reaches 0.6. IPTG was then added to the cells at a final concentration of 0.5 mM to induce the protein expression at 16 °C overnight. Cells were harvested by centrifugation and lysed by sonication in PBS lysis buffer (140 mM NaCl, 2.7 mM KCl, 10 mM Na2HPO4, 1.8 mM KH2PO4, adjusted to pH 7.4). Precleared cell lysis was then subjected for protein purification using a gravity flow column packed with Glutathione Sepharose 4B (GE Healthcare Bio-Sciences, Pittsburgh, PA) according to the manufacturer’s instructions. Fractions of eluted protein were analyzed by SDS gel electrophoresis and Coomassie blue staining, and further concentrated to a final concentration of ~1μg/μl using protein concentrators (ThermoFisher Scientific, Waltham, MA),

#### HEN1 methyltransferase activity assays

RNA oligo concentrations were quantified by Qubit Fluorometer (Invitrogen, Carlsbad, CA). Equal molarity of RNA oligos were annealed in duplex buffer (100 mM Potassium Acetate, 30 mM HEPES, pH 7.5) by heating up to 94 °C for 2 minutes, followed by cooling at room temperature. Each 50 ⍰l reaction contains 0.1 nmol of RNA duplexes (or single stranded RNA oligos), 2 μg GST-HEN1, 5 ul NEB buffer II (10X), 20 U SUPERase·In RNase Inhibitor (Invitrogen, Carlsbad, CA), and 0.5 μl SAM (10mM stock solution). Reactions were incubated at 37 °C and terminated by heating up to 70 °C for 5 minutes.

#### LC-MS/MS

RNA oligos from reactions of HEN1 methyltransferase activity assays were purified using Oligo Clean & Concentrator kit (Zymo Research, Irvine, CA), and digested by 1 U nuclease P1 from *Penicillium citrinum* (Sigma-Aldrich, St. Louis, MO) in digestion buffer containing 25 mM NaCl and 2.5 mM ZnCh at 37 °C for 2 hours, followed by dephosphorylation using 1 U FastAP Thermosensitive Alkaline Phosphatase with supplied FastAP buffer (ThermoFisher Scientific, Waltham, MA) at 37°C for 1 hour. The reactions were then adjusted to 100 μl and filtered through Millex 0.22 μm PVDF filters (Millipore, Burlington, MA) before LC-MS/MS. Ten μl of each sample was injected to a C18 reversed phase HPLC column (ThermoFisher Scientific, Waltham, MA) for compound separation, followed by mass spectrometry detection using Agilent 6410 QQQ triple-quadruple LC mass spectrometer. The nucleosides were quantified using the nucleoside-to-base ion mass transitions of 258 to 112 (Cm), 282 to 136 (Am), and 284 to 152 (G). Quantification was achieved by the comparison with the standard curves obtained from nucleoside standards injected in the same batch. The ratio of Cm or Am to G was calculated based on the standard curve-derived concentrations, and was averaged from three individual HPLC injections.

#### Accessions

Accessions of published datasets used in this study are GSE58779 (soybean) and GSE67409 (common bean) from Gene Expression Omnibus (GEO). The small RNA sequencing data generated in this study will be available at GEO.

**Fig. S1.**
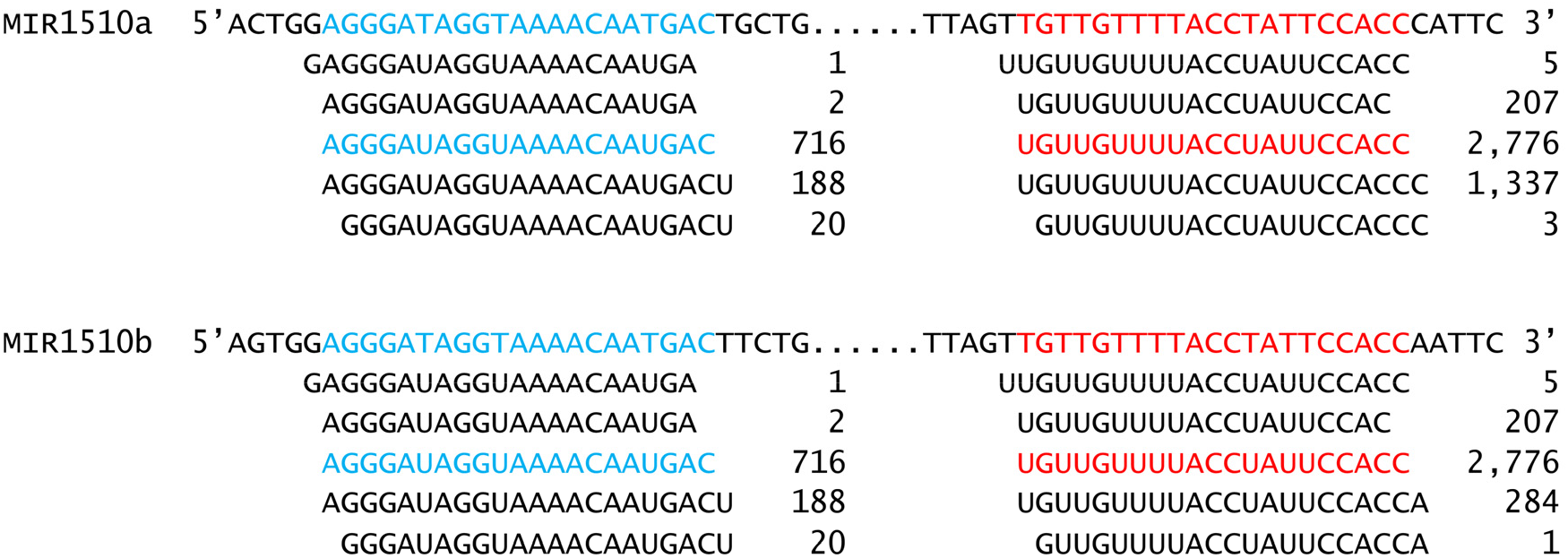
Endogenous processing of miR1510 from pri-miR1510 transcripts. Small RNA reads from the same leaf sample as in Figure 1a. Small RNA reads are normalized to reads per 10 million reads (RP10M). At top is the precursor sequence for comparison, with the mature miR1510a indicated in red and miR1510* in blue. Small RNA abundances are shown on the right of each sequence. Due to the sequence similarity between *MIR1510a* and *MIR1510b*, the same abundances are indicated for reads mapped to both *MIR1510a* and *MIR1510b*.

**Fig. S2.**
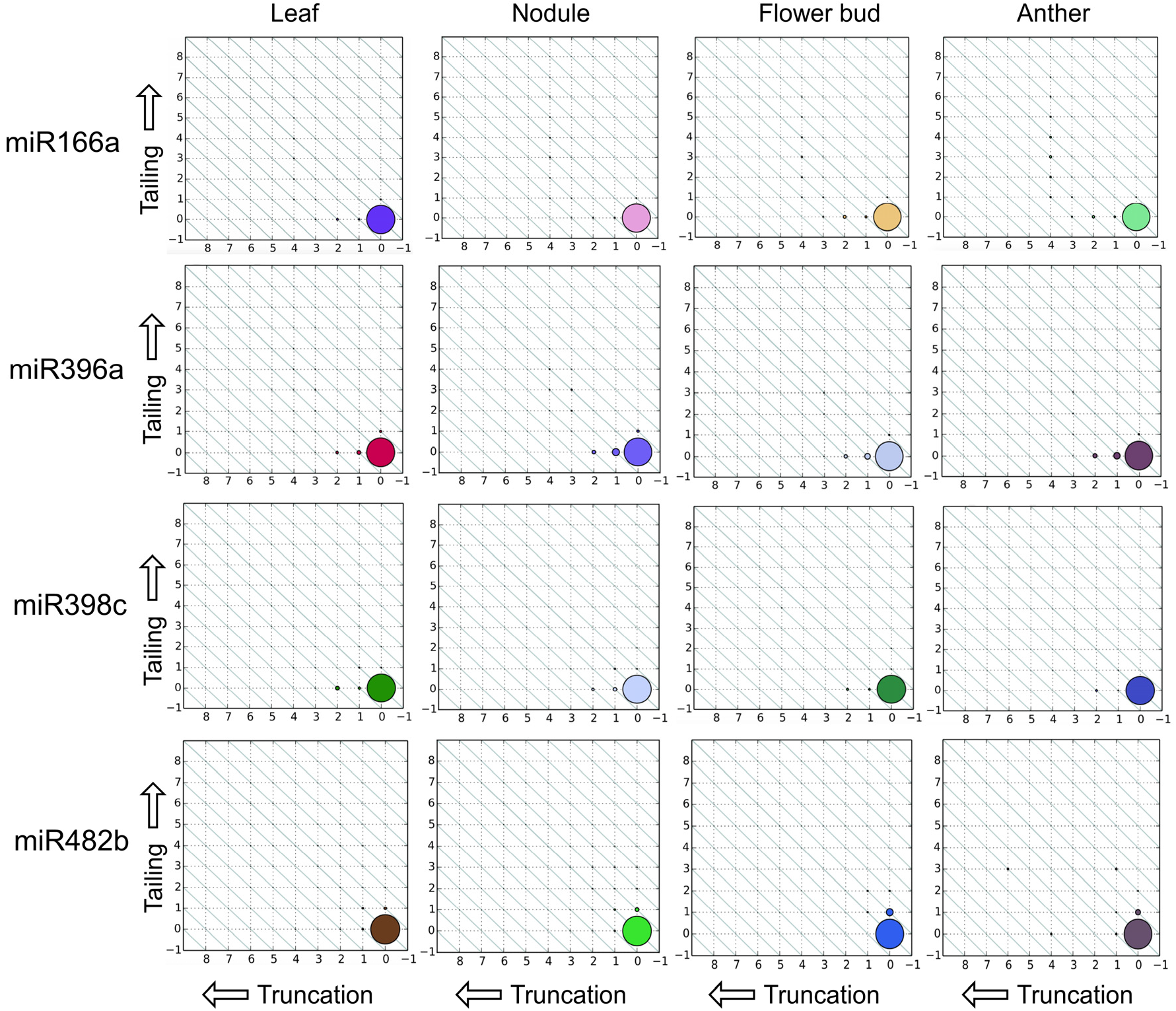
Truncation and tailing analysis for selected, exemplar miRNAs in wildtype soybean. Selected miRNAs were analyzed to represent highly conserved miRNAs, for comparison to the data for miR1510, which is shown in Figure 1. The same datasets were used for analysis as in Figure 1, and the plot matrix representation is as described in the legend of Figure 1.

**Fig. S3.**
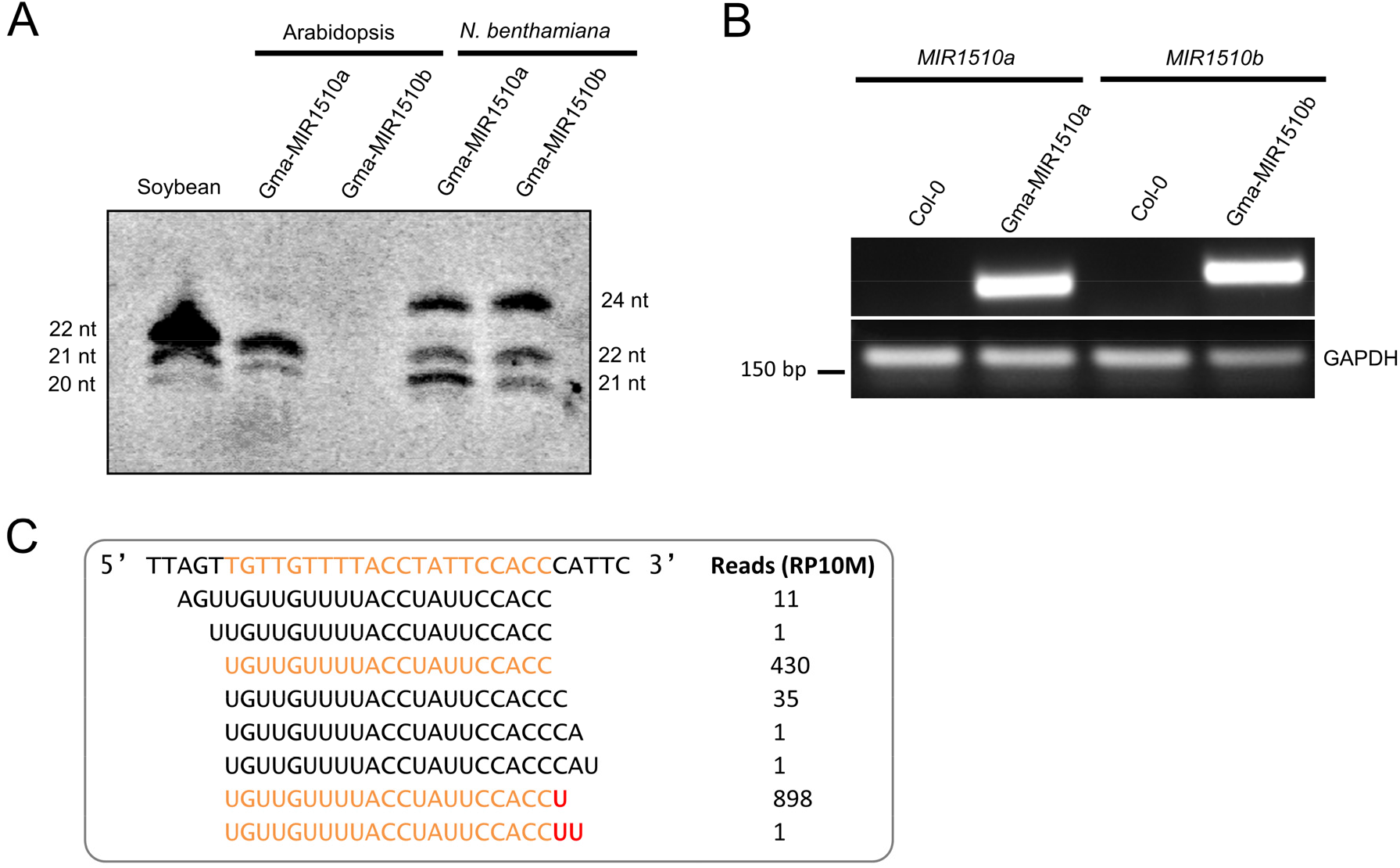
Ectopic expression of soybean miR1510 in Arabidopsis and *N. benthamiana*. A. The length of miR1510 examined by RNA gel blotting, in wildtype soybean compared to transgenes expressed in transformed Arabidopsis and *N. benthamiana*. RNA was extracted from the leaf of Arabidopsis *35S:MIR1510a/b* stable transgenic lines and Agrobacteria-infiltrated leaf of *N. benthamiana* (transient expression). The 24 nt band is generated because of transgene silencing. B. RT-PCR confirms the expression of soybean *MIR1510a* and *MIR1510b* in transgenic Arabidopsis. Precursors of *MIR1510a* or *MIR1510b* (both derived from soybean) were amplified by PCR (18 cycles) using cDNA derived from RNA extracted from transgenic Arabidopsis plants. Primers used for *MIR1510a* and *miR1510b* are listed in Table S2. C. Small RNA sequencing results confirm the uridylation of transgene-produced miR1510 in Arabidopsis transformed with the soybean *MIR1510a* precursor. Small RNA reads are normalized to reads per 10 million reads (RP10M). At top is the precursor sequence for comparison, with the mature miR1510a indicated in orange; below sequences precisely matching miR1510a or plus post-processing uridylation (in red) are indicated in orange. The sequencing data in this figure together with the soybean sequencing data from Figure 1 confirm the band sizes of miR1510 in wildtype soybean and transgenic Arabidopsis in panel A.

**Fig. S4.**
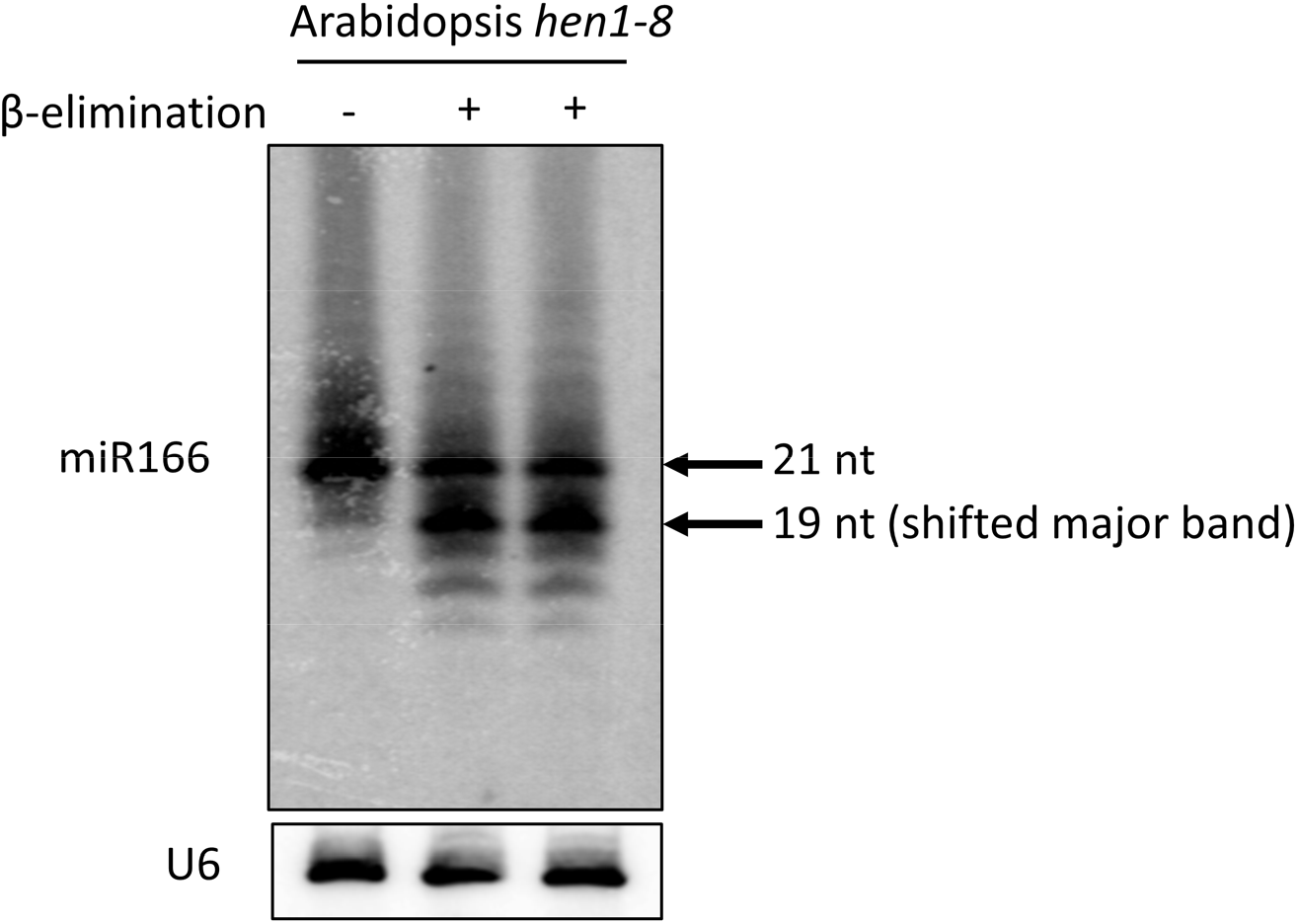
The 3’-terminal 2’-O-methylation status of Arabidopsis *hen1-8* miR166 assessed by β-elimination. Untreated RNA is loaded on the left (−), while two replicate, treated samples are on the right (+). Due to incomplete methylation in the *hen1* mutant background, miR166 is largely shifted to 19 nt after β-elimination.

**Fig. S5.**
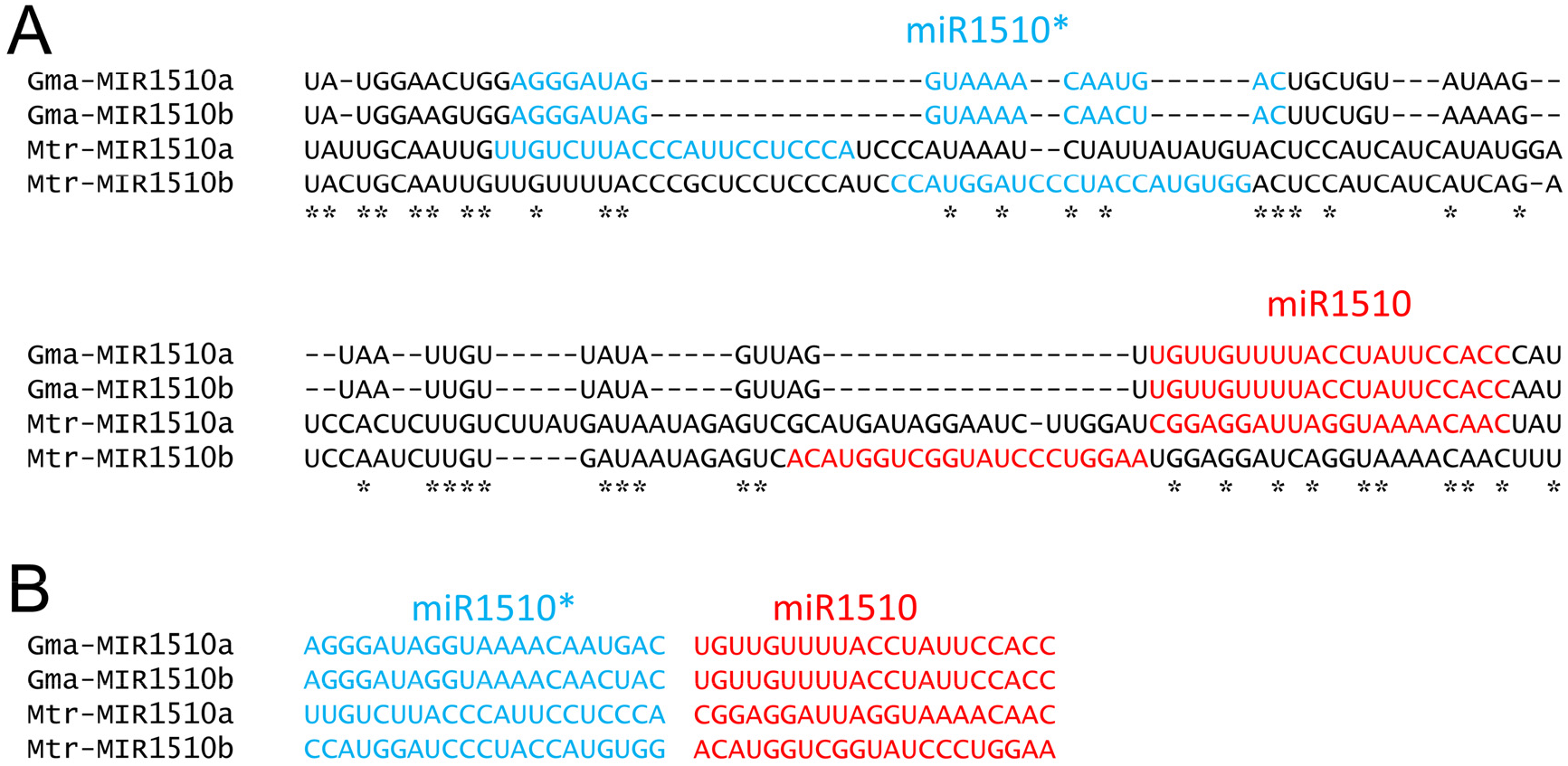
Sequence alignment reveals different origins of Medicago *MIR1510* genes. A. *MIR1510a* and *MIR1510b* precursor sequences from soybean (“Gma-”) and *Medicago truncatula* (“Mtr-”) were aligned, demonstrating a high degree of divergence. The *M. truncatula MIR1510a/b* sequences are from miRBase. The multiple sequence alignment was performed using MUSCLE. Consensus nucleotides are indicated by asterisks below the alignment. B. For clarity, only the mature small RNA sequences are shown, extracted from the alignment in panel A.

**Fig. S6.**
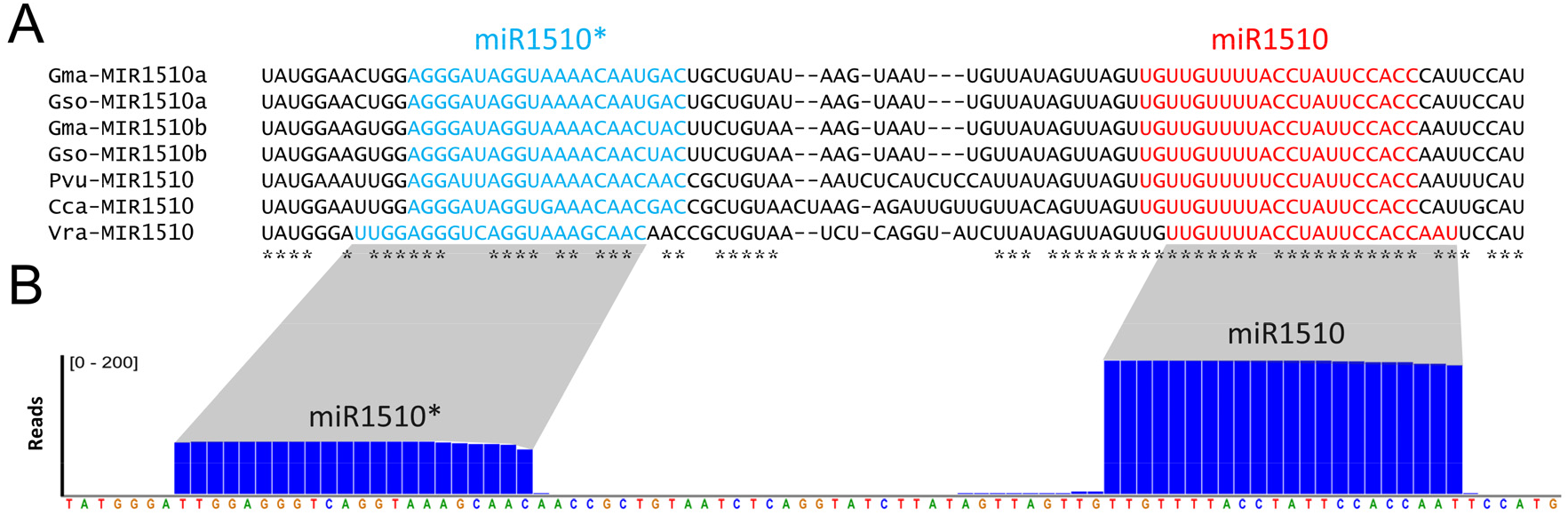
The processing of mung bean miR1510 by DCL1 is different from other species. A. Sequence alignment of *MIR1510* sequences in different species, with the mature miRNA and miRNA* sequences in red and blue, respectively, determined as the most abundant reads in the small RNA sequencing data. Multiple sequence alignment was performed using MUSCLE. Consensus nucleotides are indicated by asterisks below. B. Small RNA reads mapped to the mung bean *MIR1510* precursor sequence. A small RNA library was prepared from the leaf tissue of two-week old mung bean plants. The shaded trapezoids indicate the correspondence to the precursor in panel A.

**Figure S7.**
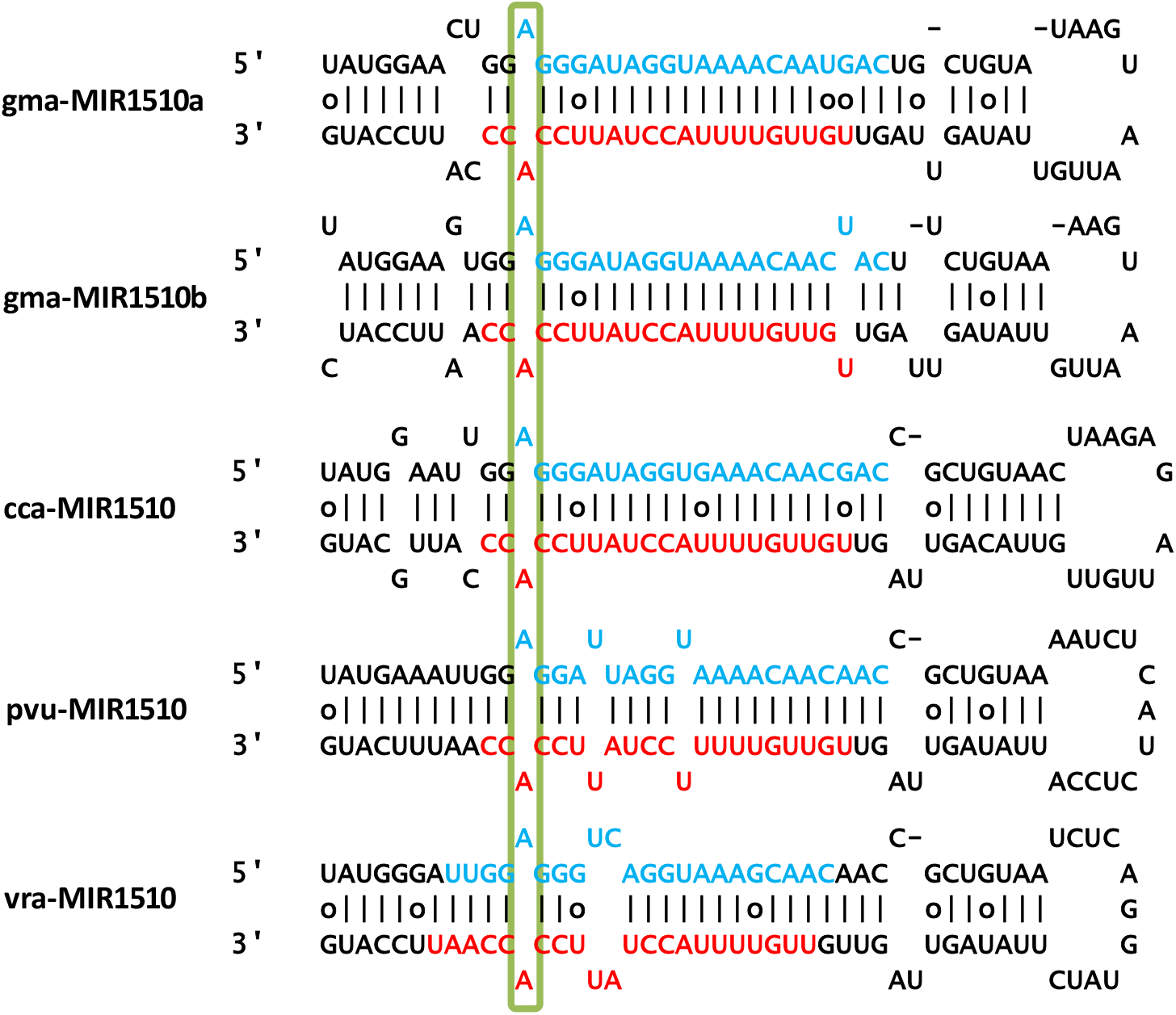
A conserved, mispaired site in the secondary structures of *MIR1510* precursors in Phaseoleae species. An unpaired set of nucleotides is predicted to exist for the miR1510/miR1510* duplex after DCL1 processing, for multiple species in the Phaseoleae tribe, indicated by the green box. Three letter species codes are as described in Figure 4. As described in the text, this mispairing is at one end of the duplex and thus may inhibit HEN1 activity to promote uridylation. Based on the predominant variants present in sequencing data from each of these species, the miRNA strand is indicated in red, while the miRNA* strand is in blue. The abundant, processed form of miR1510 from mung bean (“Vra”) is shifted by a few nucleotides, resulting in direction production of a 22-nt miRNA with no requirement for uridylation (shown in more detail in Figure S6).

**Figure S8.**
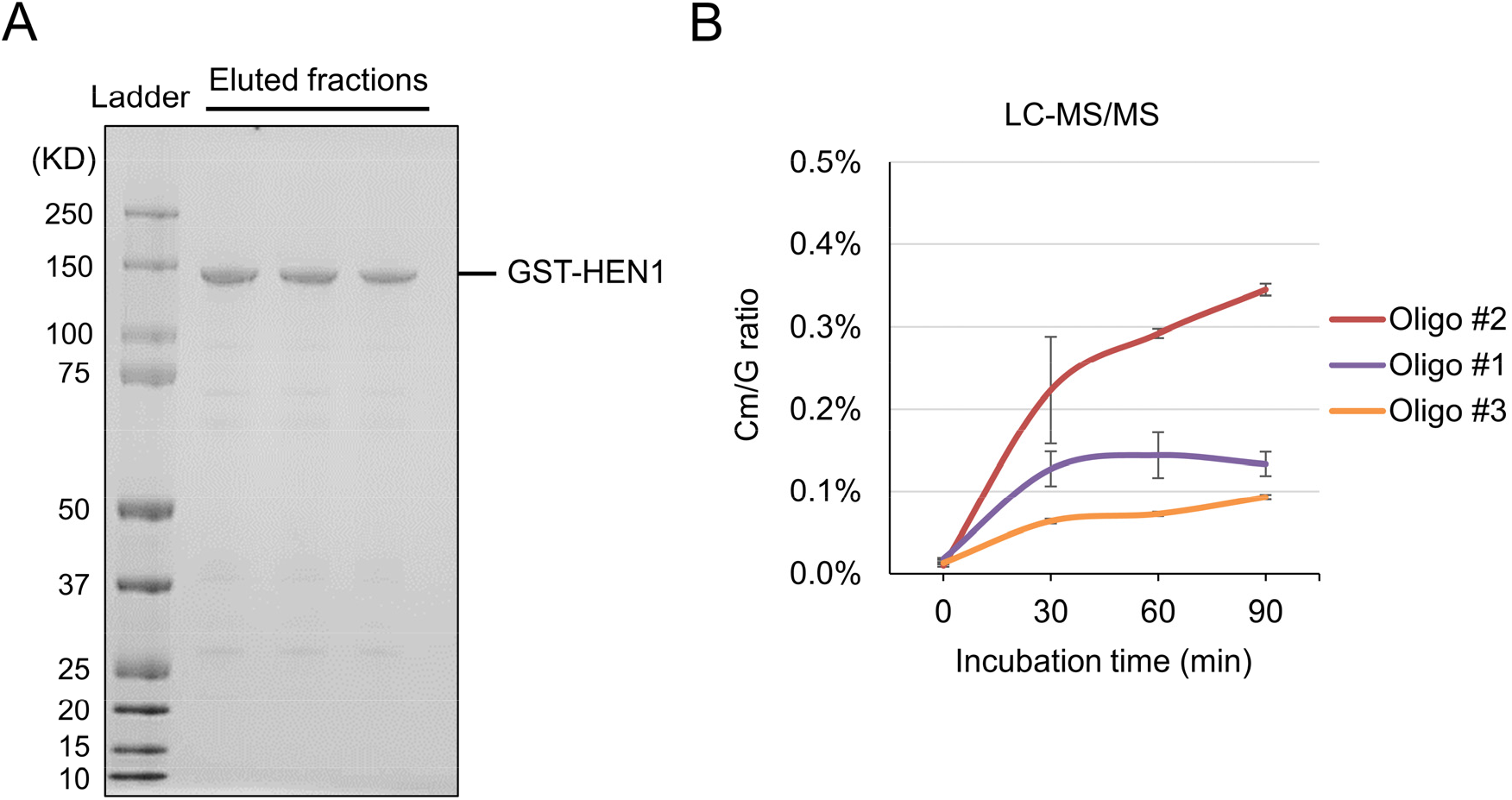
GST-HEN1 protein purification and methyltransferase activity assays. A. GST-HEN1 was purified using the Glutathione Sepharose 4B column. Different fractions of eluted proteins were analyzed by SDS gel electrophoresis followed by Coomassie blue staining. The GST-HEN1 recombinant protein is about 130 KD, as indicated as the major band in the gel image. B. HEN1 methyltransferase activity assay on single stranded RNA oligos. Oligo sequences are as indicated in Figure 5A.

**Table S1.**
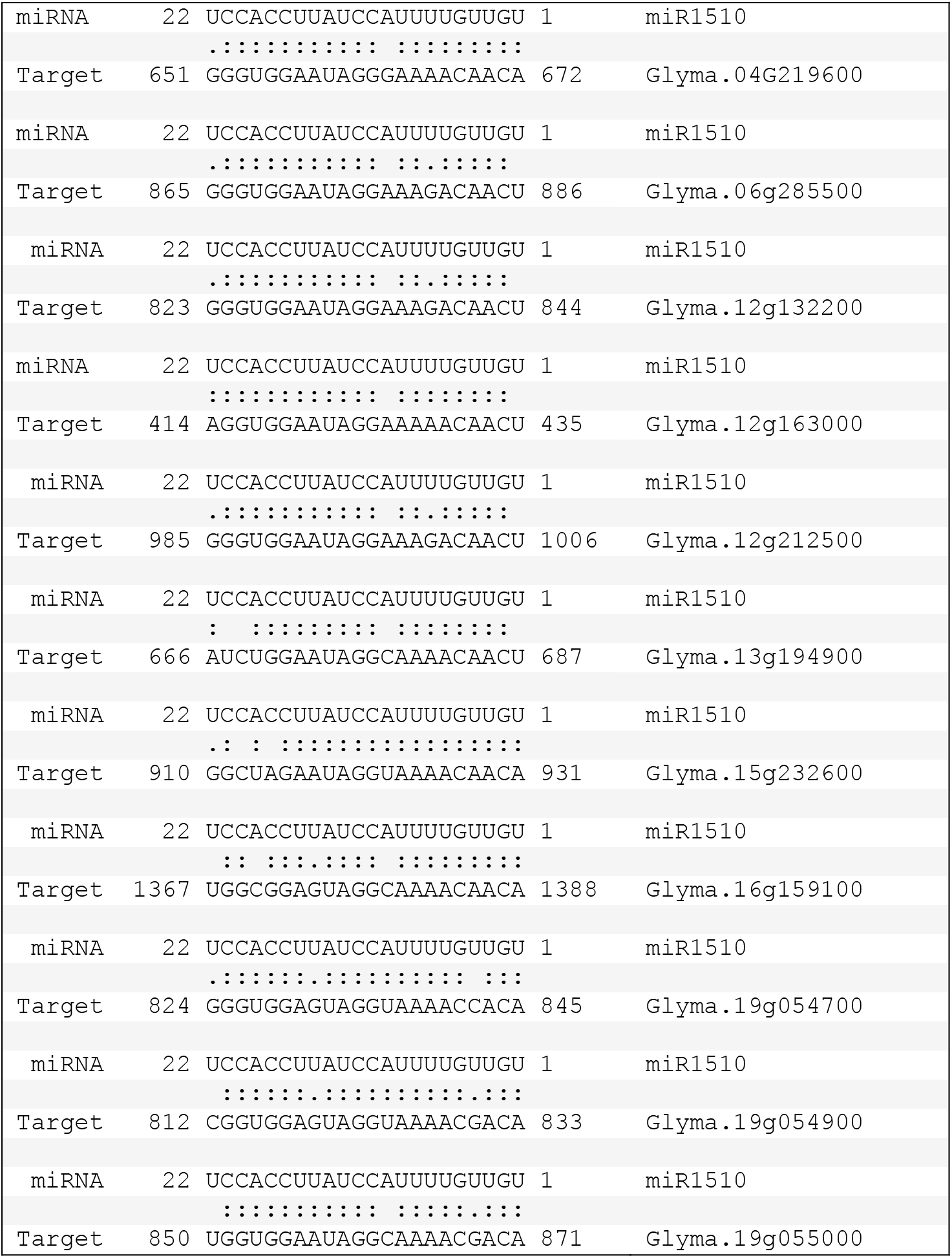
Target sites of miR1510 in *NB-LRRs* that produce phasiRNAs in soybean.

**Table S2.**
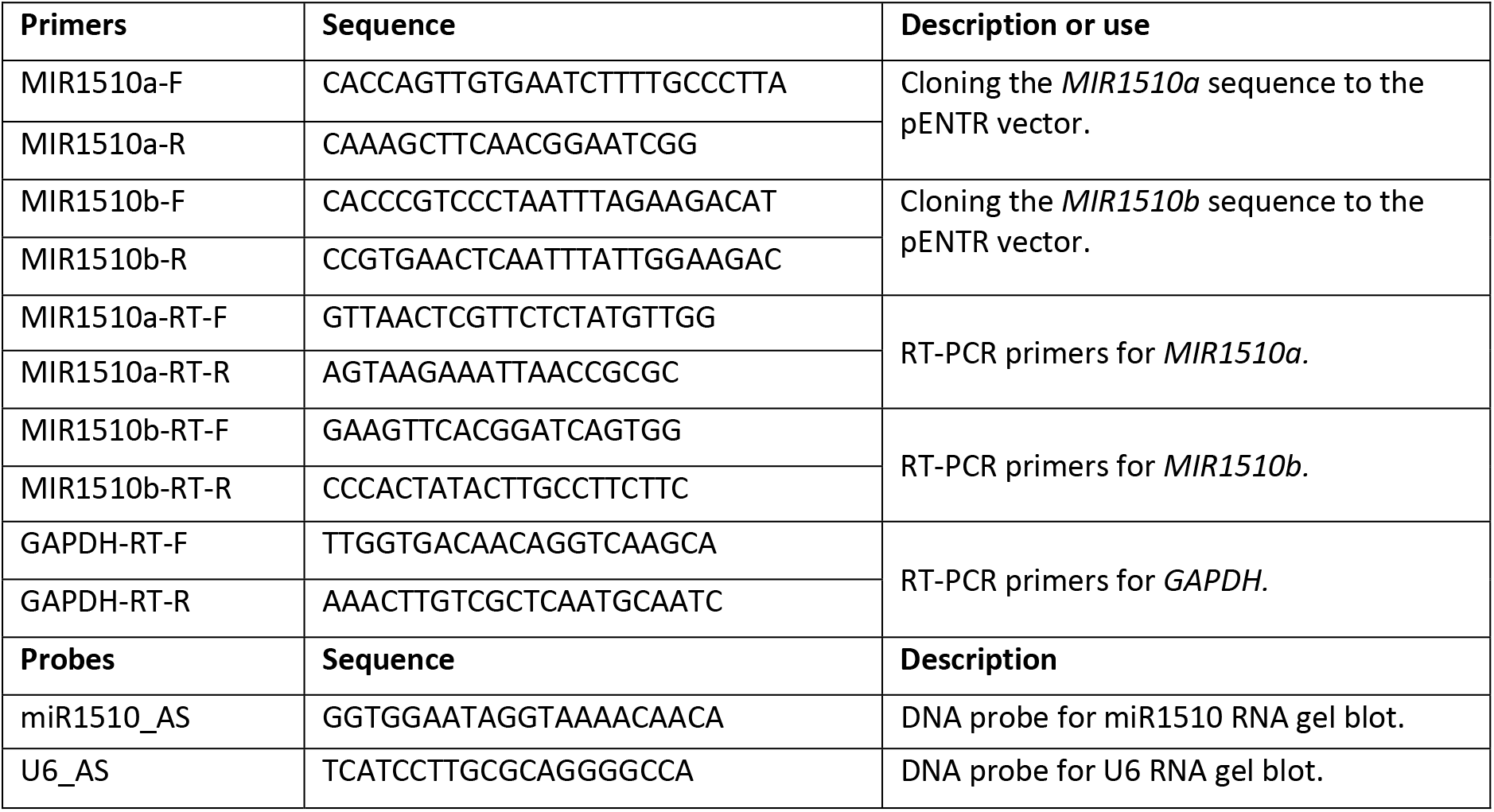
Primers and probes used in this study.

